# Recovery of infectious Oz virus from cloned cDNA

**DOI:** 10.1101/2025.11.12.687958

**Authors:** Tofazzal Md Rakib, Lipi Akter, Hiroshi Shimoda, Yudai Kuroda, Hiromichi Matsugo, Shuzo Urata, Yusuke Matsumoto

**Affiliations:** Transboundary Animal Diseases Research Center, Joint Faculty of Veterinary Medicine, Kagoshima University, Kagoshima, Japan; Department of Pathology and Parasitology, Faculty of Veterinary Medicine, Chattogram Veterinary and Animal Sciences University, Chattogram, Bangladesh; Laboratory of Veterinary Microbiology, Joint Faculty of Veterinary Medicine, Yamaguchi University, Yamaguchi, Japan; Department of Veterinary Science, National Institute of Infectious Diseases, Japan Institute for Health Security, Tokyo, Japan; Laboratory of Veterinary Public Health, Graduate School of Agricultural and Life Sciences, University of Tokyo, Tokyo, Japan; National Research Center for the Control and Prevention of Infectious Diseases (CCPID), Nagasaki University, Nagasaki, Japan

**Keywords:** Oz virus, Thogotovirus, reverse genetics

## Abstract

Oz virus (OZV) is a tick-borne, six-segmented, negative-strand RNA virus in the genus *Thogotovirus*, family *Orthomyxoviridae*. A fatal human infection was reported in Japan in 2023. In this study, we established a reverse genetics system to generate infectious recombinant OZV. Six plasmids encoding the full-length OZV genome segments under a murine RNA polymerase I promoter, together with four plasmids expressing viral proteins essential for polymerase activity, were co-transfected into murine cells. This approach enabled efficient recovery of infectious OZV. The recovered recombinant virus exhibited replication kinetics comparable to wild-type OZV. This system provides a platform for molecular studies of OZV.

## Main text

Oz virus (OZV) is a recently identified member of the genus *Thogotovirus* in the family *Orthomyxoviridae*. Members of this genus are mainly transmitted by various species of hard and soft ticks (1). OZV was first isolated in 2018 from a pool of three *Amblyomma testudinarium* nymphs collected in Ehime Prefecture, Japan (2). To date, only one human case of OZV infection has been reported in Japan, in which the patient developed fatigue, loss of appetite, vomiting, joint pain, and fever, and subsequently died due to viral myocarditis confirmed by postmortem clinical and pathological analyses (3). These findings indicate that OZV represents a potential emerging tick-borne pathogen of medical importance.

Viruses belonging to the genus *Thogotovirus* possess a six-segmented, negative-sense single-stranded RNA genome. The negative-sense RNA genome is encapsidated by nucleoprotein (NP), enabling its recognition as a template by the viral RNA-dependent RNA polymerase (RdRp) (4). The RdRp complex consists of PA, PB1, and PB2 proteins and is responsible for both genome replication and mRNA transcription (5). However, the detailed replication mechanism of OZV remains largely unknown. Reverse genetics systems that allow the recovery of infectious negative-strand RNA viruses including viruses in the genus *Thogotovirus* from cloned cDNAs are powerful experimental tools used to analyze viral replication and pathogenicity (6,7). The establishment of such a virus recovery system for OZV will provide a foundation for studies investigating its molecular biology and for future research aimed at developing antiviral strategies.

Full-length genome cDNAs of the OZV EH8 strain (GenBank accession numbers: Segment 1, NC_040730.1; Segment 2, NC_040731.1; Segment 3, NC_040732.1; Segment 4, NC_040735.1; Segment 5, LC320127.2; and Segment 6, LC320128.2) were obtained as synthetic DNA generated by GenScript’s custom gene synthesis services (GenScript Japan, Tokyo, Japan). Each cDNA segment was cloned between the murine RNA polymerase I promoter and terminator in the pRF42 vector (8), and the transcript expressed as a positive sense RNA (Fig. 1a). In addition, expression plasmids encoding OZV NP, PA, PB1, and PB2 proteins were generated using the pCAGGS vector as previously described (9). All ten plasmids were transfected into BHK/T7-9 cells seeded in 6-well plates (1×10^5^ cells/mL), using XtremeGENE HP DNA Transfection Reagent (Merck, Darmstadt, Germany), at final amounts of 0.5 μg for each full-genome plasmid and 0.4 μg for each NP, PA, PB1, and PB2 expression plasmid (Fig. 1b). As a negative control, Segment 1 and Segment 2 were replaced with an empty pCAGGS vector.

**Figure 1.**
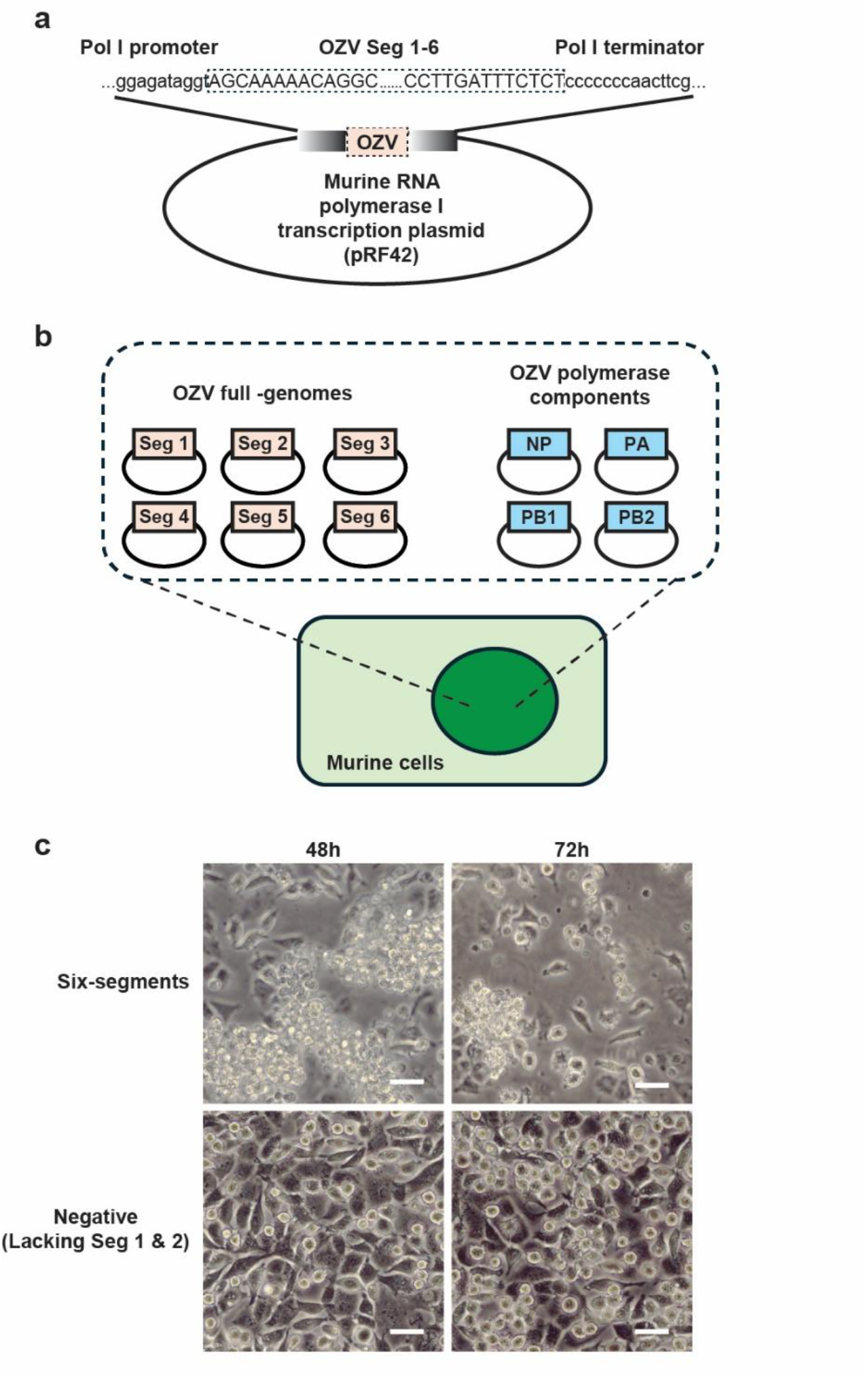
Establishment of OZV recovery system. **(a)** Schematic representation of the pRF42 plasmid containing the full-length positive-sense OZV genome inserted between the RNA polymerase I promoter and terminator. **(b)** pRF42 plasmids for OZV segments 1–6 together with pCAGGS expression plasmids for NP, PA, PB1, and PB2 were transfected into BHK/T7-9 cells. **(c)** Cytopathic effects in L929 cells observed 48 and 72 hours after co-cultivation with BHK/T7-9 cells transfected with all six segments or with segment 1 and 4 omitted. Scale bars represent 50μm.

BHK/T7-9 cells (10) were used because our previous study demonstrated that the OZV polymerase complex exhibited highly efficient activity in a minigenome assay when expressed in this cell line (9). We also confirmed that although BHK/T7-9 cells support viral replication, they do not exhibit apparent cytopathic effects (CPE). Therefore, to monitor CPE caused by OZV, transfected BHK/T7-9 cells were co-cultured with L929 mouse fibroblast cells. In the condition where all six genome segments were introduced, cell death of L929 cells was observed at 48 hours post-co-culture, and most cells were dead at 72 hours (Fig. 1c). In contrast, no CPE was observed when Segment 1 and Segment 2 were omitted (Fig. 1c). These results indicated that infectious OZV was successfully generated.

After freeze-thawing the cells, the supernatant obtained after removing cell debris by centrifugation was used to infect Vero cells, which were then incubated for 72 hours to prepare stock virus. Viral titers determined by plaque assay using Vero cells were 1.55×10^7^, 1.28×10^7^, and 1.64×10^7^ PFU/mL for recombinant OZV (rOZV) #1, #2, and #3, respectively, in three independent experiments. Furthermore, these three rOZV preparations, together with wild-type OZV EH8 strain that is gifted by National Institute of Infectious Diseases, Japan Institute for Health Security, Tokyo, Japan, were inoculated into Vero cells at a multiplicity of infection (MOI) of 1. Supernatants were harvested at 24, 48, and 72 hours post-infection and subjected to plaque assays using Vero cells. The recombinant viruses exhibited nearly identical growth kinetics to the wild-type virus (Fig. 2).

**Figure 2.**
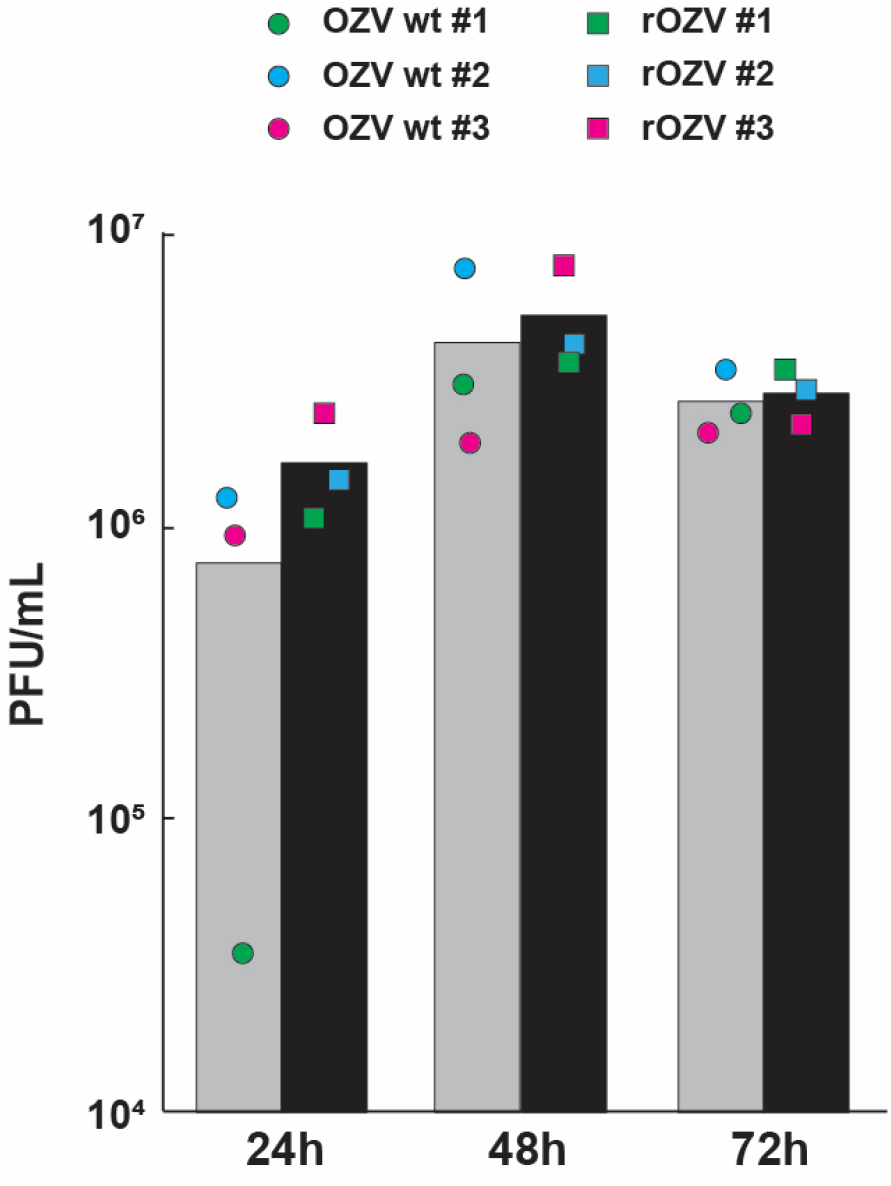
Comparison of growth kinetics between wild-type and recombinant OZV. Vero cells were infected with the wild-type OZV (OZV wt, EH8 strain) or three plasmid-derived recombinant OZVs (rOZV #1–3) at an MOI of 1. Virus titers in the supernatants at 24, 48, and 72 hours post-infection were determined by plaque assay using Vero cells. Data for OZV-wt represent the mean of three independent experiments, whereas data for rOZV represent the mean of three independent clones generated by reverse genetics. The gray bars indicate the mean titers of OZV-wt, and the black bars indicate the mean titers of rOZV. No significant differences were observed between OZV wt and rOZV at any time point (t-test, *p* > 0.05).

We also performed experiments in which plasmids were directly transfected into L929 cells without using BHK/T7-9 cells. In this case, severe cytotoxicity induced by the transfection was observed in L929 cells, resulting in extremely low viral recovery efficiency. Therefore, the method employing BHK/T7-9 cells was shown to be more efficient. In addition, when pCAGGS NP, PA, PB1, and PB2 were omitted and only the six genome-segment plasmids were transfected, no CPE was observed, indicating that the helper plasmids are essential for infectious virus production.

In summary, we successfully established a synthetic system to generate infectious OZV. This system enables the production of recombinant OZV with various mutations or modifications and will contribute to advancing studies on viral replication mechanisms and pathogenicity.

## Acknowledgements

We thank Dr. Naoto Ito (Gifu University) for providing the BHK/T7-9 cells. We also thank Dr. Haruhiko Isawa (National Institute of Infectious Diseases, Japan Institute for Health Security) for providing the OZV EH8 strain. L929 cells were obtained from RIKEN Cell Bank (RIKEN BioResource Research Center).

## Ethic Statement

The authors have nothing to report.

## Data Availability Statement

The authors have nothing to report.

